# Investigating the Association Between Body Fat and Depression via Mendelian Randomization

**DOI:** 10.1101/539601

**Authors:** Maria S. Speed, Oskar H. Jefsen, Anders D. Børglum, Doug Speed, Søren D. Østergaard

## Abstract

Obesity and depression are major public health concerns that are both associated with substantial morbidity and mortality. There is a considerable body of literature linking obesity to the development of depression. Recent studies using Mendelian Randomization indicate that this relationship is causal. Most studies of the obesity-depression association have used body mass index as a measure of obesity. Body mass index is defined as weight (measured in kilograms) divided by the square of height (meters) and therefore does not distinguish between the contributions of fat and non-fat to body weight. To better understand the obesity-depression association, we conduct a Mendelian Randomization study of the relationship between fat mass, non-fat mass, height, and depression, using genome-wide association study results from the UK Biobank (*n*=332,000) and the Psychiatric Genomics Consortium (*n*=480,000). Our findings suggest that both fat mass and height (short stature) are causal risk factors for depression, while non-fat mass is not. These results represent important new knowledge on the role of anthropometric measures in the etiology of depression. They also suggest that reducing fat mass will decrease the risk of depression, which lends further support to public health measures aimed at reducing the obesity epidemic.

## Introduction

Obesity, defined as abnormal or excessive accumulation of body fat, is a major public health concern^1^ as an established risk factor for cardiovascular disease, type II diabetes, certain cancers, and overall decreased life expectancy.^2^ Furthermore, observational studies have shown an association between obesity and depression. For example, Luppino et al.^3^ found that obese individuals were 55% more likely to develop depression, while depressed individuals were 58% more likely to become obese.

To investigate the observed association between obesity and depression, prior studies have performed Mendelian Randomization (MR), a method from genetic epidemiology which uses data from genome-wide association studies (GWAS) to determine whether a risk factor is causal for an outcome^4^. MR studies have indicated that there is a unidirectional causal relationship going from obesity to depression, but not vice versa.^5-8^ These MR studies measured obesity using body mass index (BMI), which is calculated as weight (measured in kilograms) divided by the square of height (meters).

Although BMI is the most common measure of obesity (for example, an obese individual is generally defined as someone with BMI ≥30), its use has been repeatedly criticized.^9-11^ Most notably, BMI does not distinguish between fat mass and non-fat mass. The distinction is very important from a physiological perspective, due to the fact that adipose tissue has very different properties compared to muscle and bone, the other major tissues contributing to body weight. In particular, adipose tissue is an endocrine organ that produces a range of adipokines and inflammatory proteins, which have been associated with negative systemic effects that may also affect the brain.^12^ Furthermore, BMI does not capture body fat location, and while upper-body and visceral fat, which are most common in men, contribute to the development of an unhealthy cardiometabolic profile,^13^ lower body fat, which is more common in women, seems to be protective of this.^14^ Additionally, how much of an individual’s BMI is due to fat, and where this fat is located, might also be of importance, as these factors can influence body dissatisfaction and social stigma, with negative psychological consequences.^15^

The aim of this study was to increase the understanding of the obesity-depression association by assessing the relationship between specific and biologically informative components of BMI (fat mass (stratified on limbs and trunk), non-fat mass (stratified on limbs and trunk) and depression via a MR study using results from large GWAS.

## Methods

In total, we consider 21 anthropometric measures: the first six are BMI, weight, height, whole-body fat percentage, whole-body fat mass and whole-body non-fat mass; the remaining 15 are fat percentage, fat mass and non-fat mass for each of trunk, right arm, left arm, right leg and left leg. Supplementary Figure 1 reports the genetic correlations between the 21 measures. We use MR to test whether each measure is a causal risk factor for depression, then to test whether depression is a causal risk factor for each measure, respectively.

In order to perform a MR analysis, we require GWAS summary statistics. For the anthropometric traits, we use genome-wide summary statistics from the Neale Lab (http://www.nealelab.is/people/), who have performed association analyses for over 2000 phenotypes from the UK Biobank (http://www.ukbiobank.ac.uk/), a population-based cohort of approximately 500,000 predominantly-British individuals; the average sample size is 331,910 (see Table 1). For depression, we use summary statistics from the most recent GWAS of major depressive disorder (MDD) by the Psychiatric Genomics Consortium (PGC), available at (https://www.med.unc.edu/pgc/results-and-downloads). The PGC provides two sets of summary statistics: from their main GWAS of 480,359 samples (135,458 cases, 344,901 controls) they report results for 10,000 of the most strongly-associated SNPs; whereas from a “sub-GWAS” of 173,005 samples (59,851 cases and 113,154 controls), they report results for all SNPs. Therefore, when testing whether depression is a causal risk factor for one of the anthropometric measures (which requires only results for significantly-associated SNPs), we use summary statistics from the main GWAS, whereas when testing whether an anthropometric measure is a causal risk factor for depression (which requires genome-wide results) we use summary statistics from the sub-GWAS.

**Table 1:**
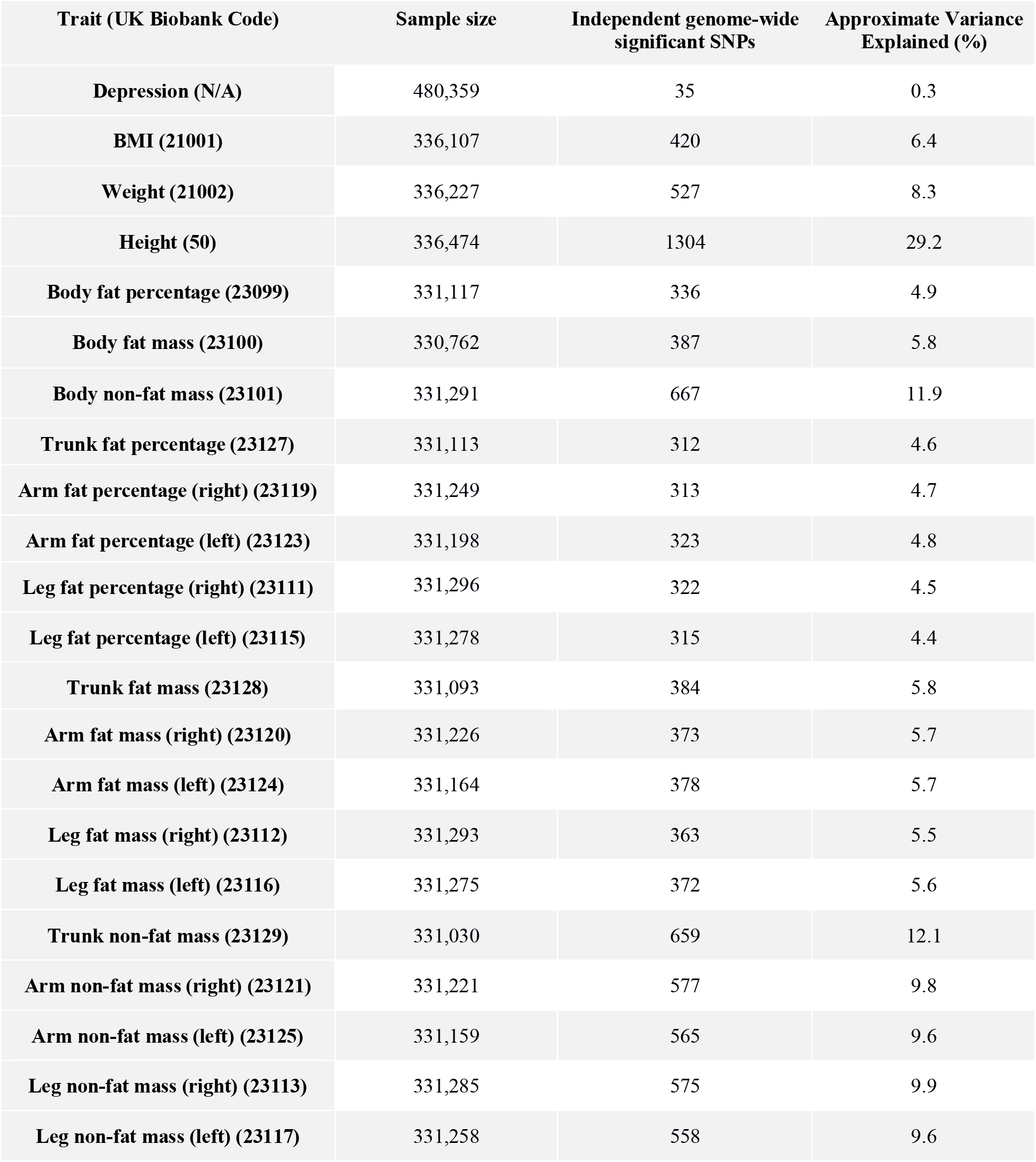
Sample size, number of independent genome-wide significant SNPs, and approximate proportion of variance these explain for depression and the 21 anthropometric measures. ^#^For depression the estimated variance is on the liability scale, assuming a prevalence of 14%.

In total, there are 6,568,396 SNPs common to the UK Biobank and PGC GWAS (we excluded SNPs with alleles A & T or C & G, or with info score<0.9). We test whether an anthropometric measure is causally associated with depression using inverse-weighted regression, for which we use the R package Mendelian-Randomization.^16^ To decide which SNPs to use in this regression, we first identify which have P<5e-8 for the measure then thin these until no pair remains within 3 centiMorgan with correlation-squared>0.05. We use the same strategy to test whether depression is a causal risk factor for an anthropometric measure, except now we identify a set of independent, genome-wide significant SNPs for depression.

A key assumption of MR is no pleiotropy.^17^ For example, the SNPs we use when testing the relationship between BMI and depression should be causal for BMI, but not depression. This is difficult to test directly, so instead we perform three sensitivity analyses. Firstly, we repeat the inverse-weighted regression excluding SNPs showing evidence for pleiotropy (P<0.05/N, where N is the number of independent, genome-wide significant SNPs). Secondly, we instead assess causality using weighted-median regression, which gives unbiased estimates provided at least 50% of the information comes from non-pleiotropic SNPs. Thirdly, we estimate the intercept from Egger Regression; an intercept significantly different to zero (P<0.05) is an indication of directional pleiotropy.

Our primary analysis focuses on BMI, weight, height, whole-body fat mass, whole-body non-fat mass and whole-body fat percentage; our secondary analysis considers the 15 location-specific measures of fat mass, non-fat mass and fat percentage. As we are performing a total of 42 tests, we set the (conservative) Bonferroni corrected significance threshold at P<0.05/42 (this is satisfied when the regression slope is ≥3.0 SDs from zero).

## Results

Table 1 reports the number of independent, genome-wide significant SNPs for each of the 21 anthropometric measures, and how much of the phenotypic variation each set of SNPs explains. The results of our primary MR analysis are displayed in Figures 1 & 2 and Table 2. Our main conclusions are based on the results from inverse-variance regression using all SNPs (the red lines in each plot), while the remaining three regressions (the orange, green and blue lines) are sensitivity analyses. We note that each time the slope from inverse-variance regression using all SNPs is statistically significant, this finding is supported by the three sensitivity analyses (i.e., the slope remains significant when SNPs showing evidence for pleiotropy are excluded or when we use weighted-median regression, and Egger regression finds no evidence for directional pleiotropy). Figure 1(a) confirms that BMI is a causal risk factor for depression. The estimated slope from inverse-weighted regression is 0.17 (SD 0.03), which is significantly greater than zero (P=1e-7) and indicates that a 1-SD increase in BMI corresponds to a 0.17 increase in the log-odds ratio (OR) for depression. Figure 1(b) indicates there is a causal relationship between weight and depression; however, the estimated slope (0.13, SD 0.03) is less than that for BMI, which supports the current preference to measure obesity using BMI rather than weight.

**Table 2:**
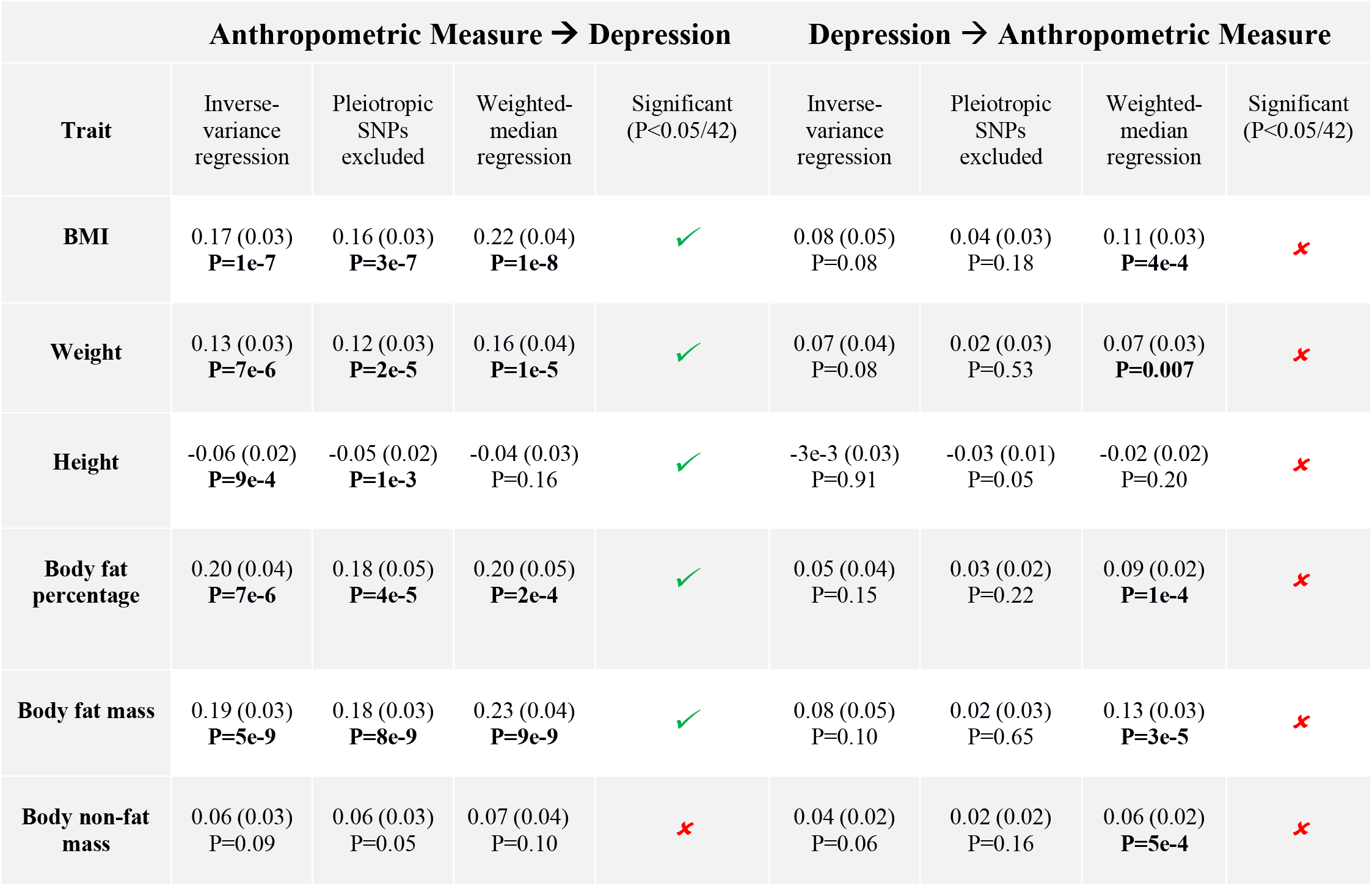
Columns 2, 3, 4, 6, 7 & 8 report estimates of the slope (SD in brackets) and p-value from inverse-weighted regression, inverse-weighted regression with pleiotropic SNPs excluded and weighted-median regression. Columns 2-5 examine whether each anthropometric measure is causal for depression; Columns 6-9 examine whether depression is causal for each anthropometric measure. Significant tests (slope estimate P<0.05/42) are marked in **bold**.

**Figure 1:**
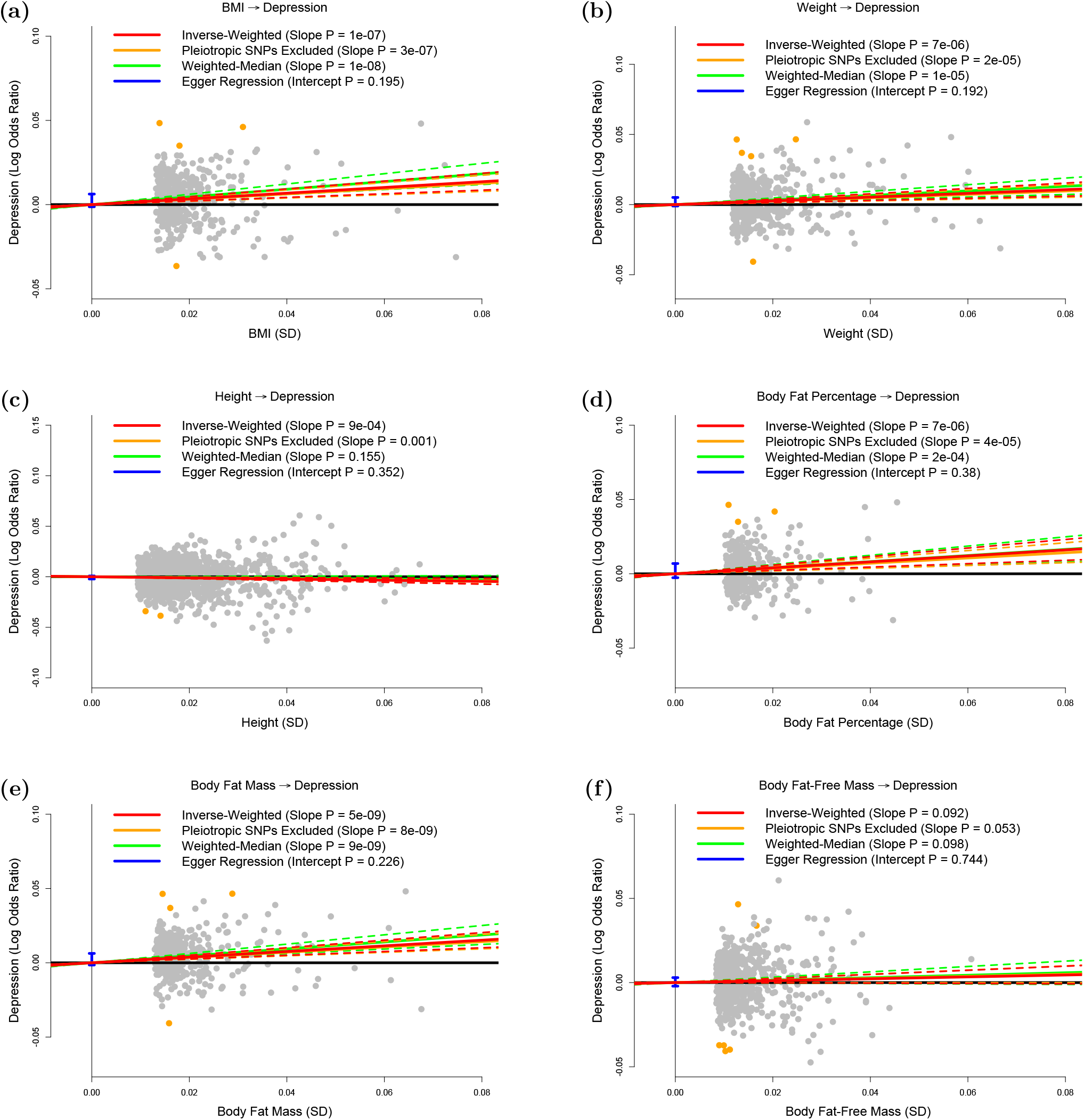
Testing whether the anthropometric measures are causal risk factors for depression. The panels plot per-allele effect sizes for **(a)** BMI, **(b)** weight, **(c)** height, **(d)** body fat percentage, **(e)** body fat mass and **(f)** body fat-free mass (x-axes) against per-allele effect size for depression (y-axis) For the anthropometric measures, effect sizes are measured in SDs, for depression the effect size is log-odds ratio. For each plot we estimate the slope using inverse-variance regression (red solid line), inverse-variance regression after excluding SNPs showing evidence for pleiotropy (orange solid line), weighted-median regression (green solid line) and Egger regression (blue solid line). The corresponding colored dashed lines represent the 95% confidence intervals for the slopes, while the vertical blue segment marks a 95% confidence interval for the intercept from Egger regression. The horizontal black solid line indicates no effect.

Figure 1(c) suggests that height (short stature) is a causal risk factor for depression; the estimated slope is −0.06 (SD 0.02), indicating that a 1-SD increase in height corresponds to a 0.06 decrease in the log-odds ratio (OR) for depression.

Figure 1(d) and 1(e) shows that whole-body fat percentage and whole-body fat mass are both causal risk factors for depression; the estimated slopes are very similar (0.20, SD 0.04 and 0.19, SD 0.03, respectively) indicating that there is no advantage to normalizing (i.e., measuring fat percentage instead of fat mass). The estimated slope in Figure 1(f) is 0.06 (SD 0.03), indicating that MR finds no evidence that whole-body non-fat is a causal risk factor for depression.

Figure 2 shows that, consistent with previous studies, there is no evidence that depression is a causal risk factor for any of the six anthropometric measures investigated here.

**Figure 2:**
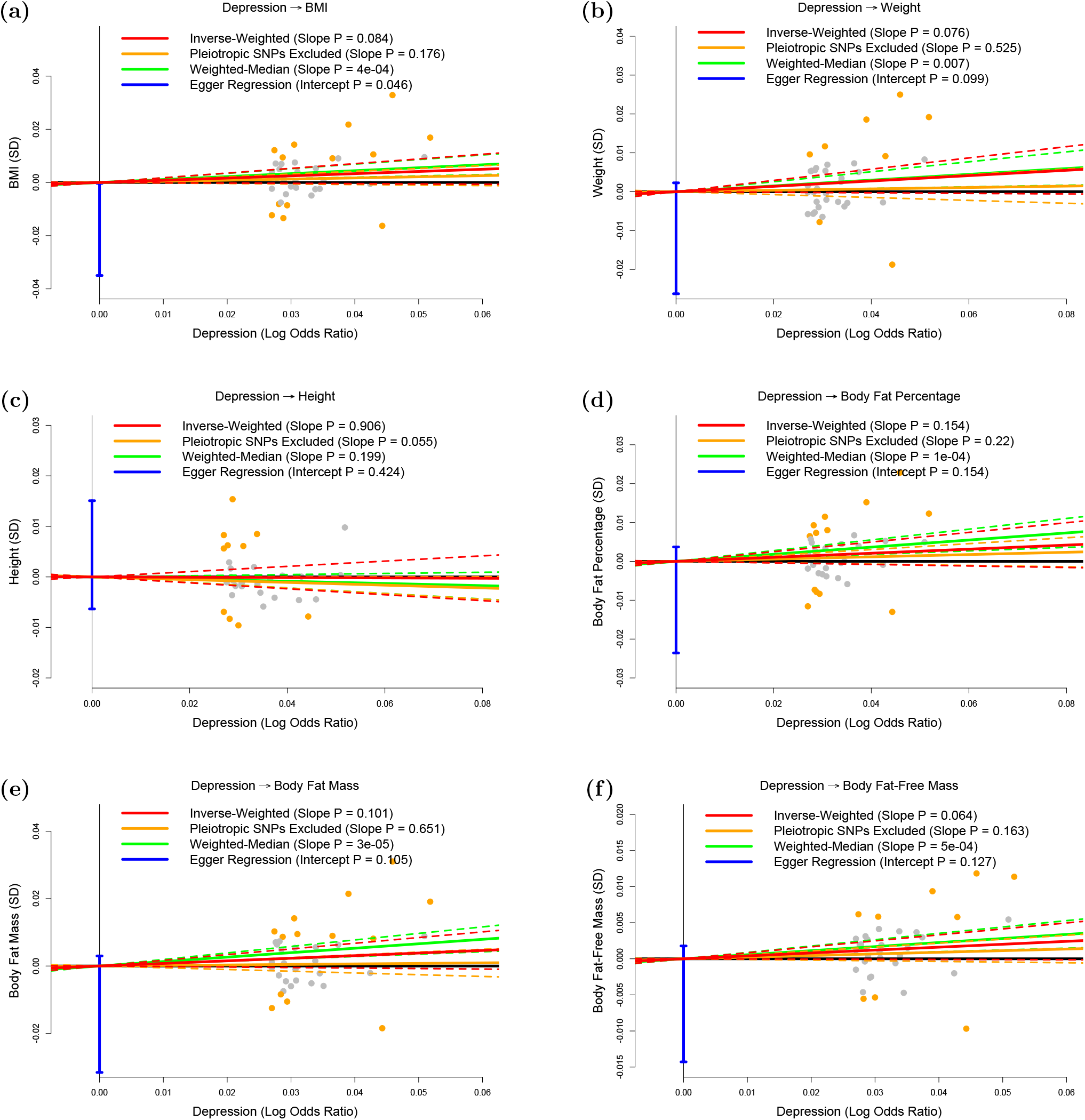
Testing whether depression is causally associated with the anthropometric measures. The panels plot per-allele effect size for depression (x-axis) against per-allele effect sizes for **(a)** BMI, **(b)** weight, **(c)** height, **(d)** body fat percentage, **(e)** body fat mass and **(f)** body fat-free mass (y-axes). For the anthropometric measures, effect sizes are measured in SDs, for depression the effect size is log-odds ratio. For each plot we estimate the slope using inverse-variance regression (red solid line), inverse-variance regression after excluding SNPs showing evidence for pleiotropy (orange solid line), weighted-median regression (green solid line) and Egger regression (blue solid line). The corresponding colored dashed lines represent the 95% confidence intervals for the slopes, while the vertical blue segment marks a 95% confidence interval for the intercept from Egger regression. The horizontal black solid line indicates no effect.

Table 3 reports results from our secondary analysis. For each of the 15 location-specific anthropometric measures, the results are consistent with those from the whole-body version; i.e., regardless of whether we consider trunk, right arm, left arm, right leg or left arm, we again find that fat percentage and fat mass are causally associated with depression, but that non-fat mass is not. For fat percentage and fat mass, we are interested in comparing slope estimates, as significant differences would indicate that risk of developing depression depends on fat location. The largest difference is observed for fat percentage, where the estimated slope for left leg (0.34, SD 0.06) is approximately twice that for trunk (0.16, SD 0.04); however, although this difference is nominally significance (P=0.01), it is not significant after correction for multiple comparisons.

**Table 3:** Columns 2, 3, 4, 6, 7 & 8 report estimates of the slope (SD in brackets) and p-value from inverse-weighted regression, inverse-weighted regression with pleiotropic SNPs excluded and weighted-median regression. Columns 2-5 examine whether each anthropometric measure is causal for depression; Columns 6-9 examine whether depression is causal for each anthropometric measure. Significant tests (slope estimate P<0.05/42) are marked in **bold**.

| Anthropometric Measure → Depression |  |  |  |  | Depression → Anthropometric Measure |  |  |  |
| --- | --- | --- | --- | --- | --- | --- | --- | --- |
| Trait | Inverse-variance<br>regression | Pleiotropic<br>SNPs<br>excluded | Weighted-median<br>regression | Significant<br>(P<0.05/42) | Inverse-variance<br>regression | Pleiotropic<br>SNPs<br>excluded | Weighted-median<br>regression | Significant<br>(P<0.05/42) |
| Trunk fat<br>percentage | 0.16 (0.04)<br>**P=3e-5** | 0.14 (0.04)<br>**P=1e-4** | 0.18 (0.05)<br>**P=1e-4** | ✓ | 0.06 (0.04)<br>P=0.16 | 0.03 (0.03)<br>P=0.34 | 0.06 (0.03)<br>P=0.04 | ✗ |
| Arm fat<br>percentage<br>(right) | 0.26 (0.05)<br>**P=5e-8** | 0.23 (0.05)<br>**P=3e-7** | 0.30 (0.06)<br>**P=3e-7** | ✓ | 0.05 (0.03)<br>P=0.11 | 0.02 (0.02)<br>P=0.32 | 0.10 (0.02)<br>**P=3e-5** | ✗ |
| Arm fat<br>percentage<br>(left) | 0.24 (0.05)<br>**P=2e-7** | 0.23 (0.05)<br>**P=9e-7** | 0.30 (0.06)<br>**P=1e-7** | ✓ | 0.06 (0.04)<br>P=0.12 | 4e-3 (0.02)<br>P=0.86 | 0.10 (0.02)<br>**P=6e-5** | ✗ |
| Leg fat<br>percentage<br>(right) | 0.33 (0.06)<br>**P=6e-9** | 0.30 (0.05)<br>**P=4e-8** | 0.36 (0.07)<br>**P=2e-7** | ✓ | 0.04 (0.03)<br>P=0.24 | 0.02 (0.02)<br>P=0.27 | 0.04 (0.02)<br>P=0.03 | ✗ |
| Leg fat<br>percentage<br>(left) | 0.34 (0.06)<br>**P=2e-8** | 0.30 (0.06)<br>**P=1e-7** | 0.38 (0.07)<br>**P=2e-7** | ✓ | 0.04 (0.03)<br>P=0.20 | 0.03 (0.02)<br>P=0.06 | 0.05 (0.02)<br>**P=7e-3** | ✗ |
| Trunk fat<br>mass | 0.17 (0.03)<br>**P=2e-8** | 0.16 (0.03)<br>**P=1e-7** | 0.20 (0.04)<br>**P=2e-7** | ✓ | 0.07 (0.05)<br>P=0.12 | -4e-3 (0.03)<br>P=0.90 | 0.09 (0.03)<br>P=0.01 | ✗ |
| Arm fat mass<br>(right) | 0.19 (0.03)<br>**P=8e-9** | 0.18 (0.03)<br>**P=2e-8** | 0.25 (0.04)<br>**P=2e-9** | ✓ | 0.08 (0.05)<br>P=0.08 | 0.02 (0.03)<br>P=0.63 | 0.14 (0.03)<br>**P=4e-6** | ✗ |
| Arm fat mass<br>(left) | 0.19 (0.03)<br>**P=2e-8** | 0.17 (0.03)<br>**P=3e-8** | 0.25 (0.04)<br>**P=5e-9** | ✓ | 0.08 (0.05)<br>P=0.08 | 0.02 (0.03)<br>P=0.47 | 0.14 (0.03)<br>**P=9e-6** | ✗ |
| Leg fat mass<br>(right) | 0.27 (0.04)<br>**P=1e-10** | 0.25 (0.04)<br>**P=2e-10** | 0.33 (0.05)<br>**P=6e-10** | ✓ | 0.06 (0.04)<br>P=0.14 | 0.03 (0.02)<br>P=0.25 | 0.09 (0.02)<br>**P=2e-4** | ✗ |
| Leg fat mass<br>(left) | 0.27 (0.04)<br>**P=1e-10** | 0.25 (0.04)<br>**P=3e-10** | 0.33 (0.05)<br>**P=2e-10** | ✓ | 0.06 (0.04)<br>P=0.13 | 0.03 (0.02)<br>P=0.22 | 0.10 (0.03)<br>**P=6e-05** | ✗ |
| Trunk non-fat mass | 0.03 (0.03)<br>P=0.28 | 0.03 (0.03)<br>P=0.28 | 0.05 (0.04)<br>P=0.27 | ✗ | 0.03 (0.02)<br>P=0.09 | 0.03 (0.02)<br>P=0.10 | 0.05 (0.02)<br>**P=2e-03** | ✗ |
| Arm non-fat<br>mass (right) | 0.04 (0.04)<br>P=0.33 | 0.04 (0.04)<br>P=0.29 | 0.03 (0.05)<br>P=0.54 | ✗ | 0.04 (0.02)<br>P=0.05 | 0.01 (0.02)<br>P=0.43 | 0.06 (0.02)<br>**P=5e-4** | ✗ |
| Arm non-fat<br>mass (left) | 0.06 (0.04)<br>P=0.13 | 0.06 (0.04)<br>P=0.11 | 0.07 (0.05)<br>P=0.15 | ✗ | 0.04 (0.02)<br>P=0.05 | 9e-3 (0.02)<br>P=0.60 | 0.05 (0.02)<br>**P=2e-3** | ✗ |
| Leg non-fat<br>mass (right) | 0.06 (0.04)<br>P=0.11 | 0.06 (0.03)<br>P=0.09 | 0.08 (0.05)<br>P=0.08 | ✗ | 0.05 (0.02)<br>P=0.05 | 0.03 (0.02)<br>P=0.13 | 0.06 (0.02)<br>**P=9e-4** | ✗ |
| Leg non-fat<br>mass (left) | 0.08 (0.04)<br>P=0.04 | 0.07 (0.04)<br>P=0.03 | 0.09 (0.05)<br>P=0.05 | ✗ | 0.05 (0.03)<br>P=0.07 | 0.03 (0.02)<br>P=0.06 | 0.05 (0.02)<br>**P=2e-3** | ✗ |

## Discussion

We have used MR to investigate the causal relationship between different anthropometric measures and depression. We first confirmed that BMI is a causal risk factor for depression, but found no evidence of the opposite, i.e., that depression causes increased BMI. These results are in line with recent MR studies reporting evidence that higher BMI causally increases the risk of depression, but not vice versa.^5-8^

Our main finding is that body fat mass is causal risk factor for depression, but that body non-fat mass is not, therefore indicating that the BMI-depression causality is driven by fat. We note from Table 1 that the number of genome-wide associated SNPs for whole-body non-fat (n=667) and the proportion of variance they explain (11.9%) are higher than the corresponding values for fat percentage (n=336 and 4.9%) and for fat mass (n=387 and 5.8%), indicating that MR finding no evidence for a causal relationship was not simply a power issue.

We also investigated whether the strength of the causal relationship between body fat and depression depended on the location, but these results were inconclusive; while the point estimates suggested that leg fat more strongly affects risk of depression than either trunk fat or arm fat, the difference was not significant after correcting for multiple comparisons.

The causal relationship going from fat mass to depression is likely to have both psychological and biological components. Psychologically, perceived weight discrimination, stigmatization, and body image dissatisfaction may mediate the causality;^18-21^ biologically, obesity is associated with several endocrine and metabolic changes that have been linked to depression, including altered glucocorticoid, adipokine, insulin, leptin, and inflammatory signaling.^22^ Although our study was not aimed at providing insight into how fat increases the risk of depression, the finding that trunk fat mass was not more strongly associated with depression risk than fat mass on the limbs (rather we found a tendency towards the opposite), seems to be in favor of a psychological mechanism – since trunk fat is considered the more metabolically adverse.^13^

Observational studies have attempted to separate the psychological and physiological components in the relationship between obesity and depression. In support of a physiological component, Jokela et al.^23^ found that the risk of depressive symptoms associated with obesity increased almost linearly with the number of metabolic risk factors; in particular, obese individuals who were metabolically unhealthy had a higher depression risk than obese individuals who were metabolically healthy (OR = 1.23; 95% CI: 1.05, 1.45). However, the same study also found that metabolically-healthy individuals who were obese had a higher depression risk than metabolically-healthy individuals who were not obese (OR = 1.29; 95% CI: 1.12, 1.50), indicating that metabolic factors only partially explain the increase in depression risk associated with obesity. Similar results were found by the longitudinal study of Hamer et al.^24^

A recent study by Tyrrell et al.^8^ also sought to separate the psychological component of obesity from its adverse metabolic consequences – and employed MR to do so. Specifically, Tyrrell et al. used two genetic instruments, that both represented BMI, but one with and one without its adverse metabolic consequences, respectively. They found that both instruments were associated with increased risk of depression suggesting that the causal association between BMI and depression is primarily driven by psychological consequences of adiposity and not by its adverse metabolic effects.^8^ This conclusion resonates well with the results of our study.

In addition to the results on the impact of body fat on the risk of depression, we also found evidence suggesting that height (short stature) is a causal risk factor for depression. Several large observational studies have found short stature to be associated with poorer mental health,^25^ lower health-related quality of life,^26^ depressive symptoms in adolescents^27^ and adults^28, 29^, and suicide in men.^30^ Other studies have found no association between height and depression or suicide^31^ or negligible effects of height on mental health.^32^ The association between short stature and poor mental health may be explained by confounding factors, such as socioeconomic status, prenatal development or childhood factors, or by a causal effect of height on depression risk.^33, 34^ Our results indicate that at least part of the association between short stature and depression is indeed due to a direct causal effect.

Finally, we note some limitations of our work. Firstly, the MR estimates rely on three key assumptions: (i) the SNPs used as genetic predictors for a trait are causal for that trait; (ii) these SNPs are not associated with confounders of the trait-outcome association; (iii) these SNPs only affect the outcome through the causal relationship (i.e., there is no pleiotropy). We can be confident of (i) because our genetic predictors only used SNPs robustly-associated (P<5e-8) with the trait, while (ii) should be true due to the fact an individual’s genotypes are randomly allocated during gamete formation. It is hard to explicitly test (iii), however, our sensitivity analyses indicate that our conclusions are not the consequence of pleiotropy.

Secondly, the UK Biobank measured fat and non-fat mass via bioelectrical impedance analysis (using a Tanita BC418MA body composition analyzer), which is considered less accurate than techniques such as dual-energy x-ray absorptiometry. However, we would expect measurement error to cause the estimates for fat and non-fat to become more similar, so the fact that we observed a significant difference indicates that the UK Biobank measurements were sufficiently accurate for our purpose.

Thirdly, while we found only suggestive evidence that location of fat affects risk of depression, we recognize that the high correlations between measurements taken in the trunk, arms and legs (Supplementary Figure 1) would have reduced our power to detect significant differences. Fourthly, it is recognized that there are sex-specific psychological and physiological factors affecting the obesity-depression relationship. While we were not able to test sex differences, because the PGC did not release results from male-only and female-only GWAS of depression, this is worthy of further study.

In conclusion, the present study provides evidence that the causal relationship between BMI and depression is driven by fat mass and height, and not by non-fat mass. These results represent important new knowledge on the role of anthropometric measures in the etiology of depression. They also suggest that reducing fat mass will decrease the risk of depression, which lends further support to public health measures aimed at reducing the obesity epidemic.

## Acknowledgements

DS is supported by the European Unions Horizon 2020 Research and Innovation Programme under the Marie Skłodowska-Curie grant agreement number 754513, by Aarhus University Research Foundation (AUFF) and the Independent Research Fund Denmark (7025-00094B). ADB is supported by grants from The Lundbeck Foundation (R102-A9118 and R155-2014-1724). Data handling and analysis on the GenomeDK HPC facility was supported by NIMH (1U01MH109514-01 to Michael O’Donovan and ADB). HPC capacity at GenomeDK was provided by the iSEQ center and Center for Genomics and Personalized Medicine, Aarhus, Denmark (grants to ADB). SDØ is supported by the Independent Research Fund Denmark (7016-00048).

## Supplementary Material

**Supplementary Figure 1:**
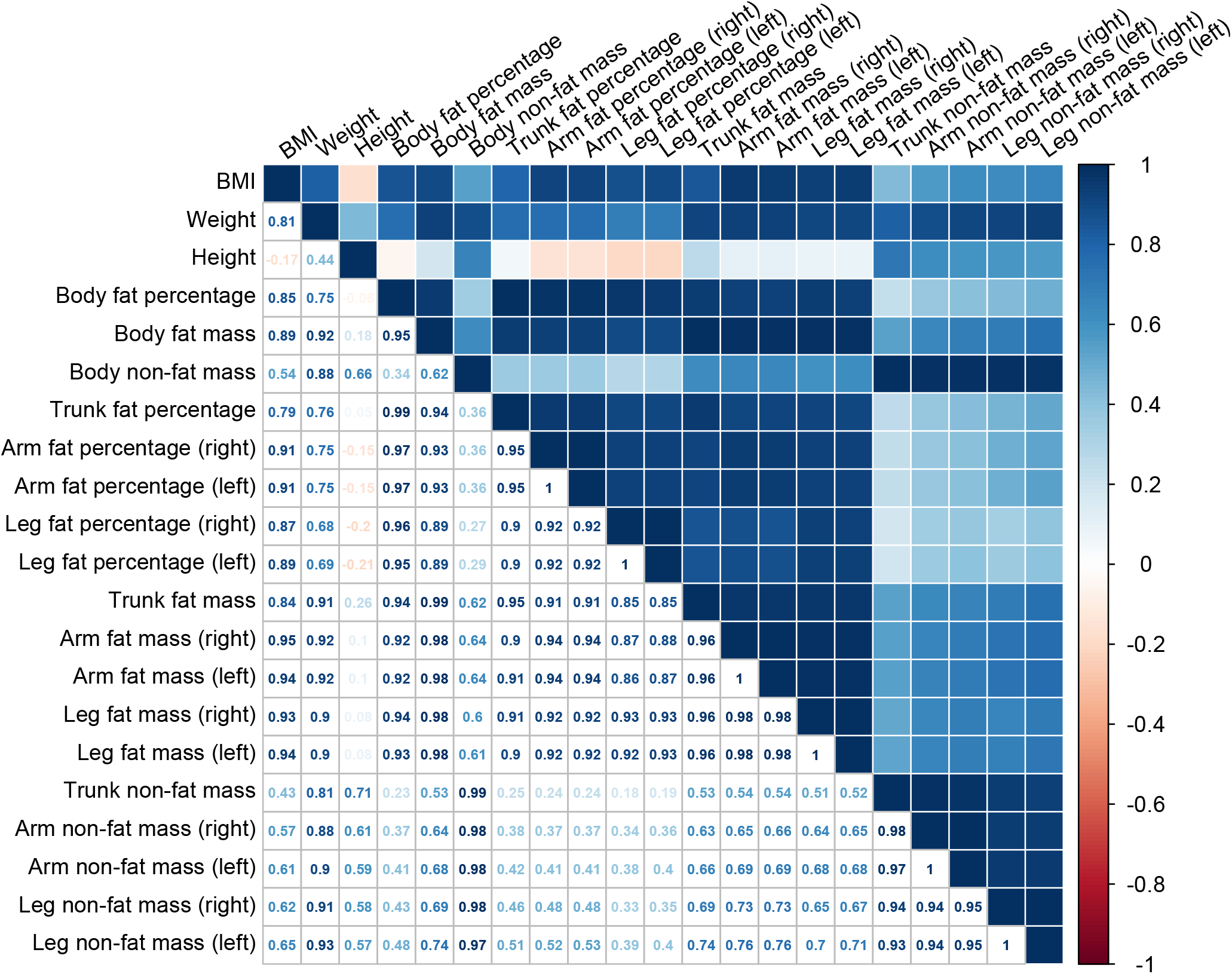
Genetic correlations. Genetic correlations between the 21 anthropometric from the UK Biobank.

